# Pathological circadian rhythm states driven by CK2 and noise

**DOI:** 10.1101/809632

**Authors:** Z. Malik, Y. Fatima, J. Alam, R. Singh

## Abstract

Circadian rhythm maintains sleep–wake cycle in living systems. Disruption of this rhythm may cause diseases. We propose an extended Drosophila circadian rhythm model incorporating cross-talk of CK2 with Per protein. We studied the model using stochastic simulation algorithm, and the behavior of the amplitude, time period and permutation entropy us identify three distinct circadian states namely, *active, weak activity, active, weak activity* and *rhythmic death* all driven by CK2. These states may correspond to distinct pathological cellular states of the living system. Noise, an important factor, has ability to switch normal circadian rhythm to any of the three aforementioned circadian states. Fluctuations in system’s size, can help us in deterning the extent of noise present. We also highlighted that disruption in circadian rhythm may lead to various diseases including cancer. We present various cellular pathways driven by per mutant genes and their pathological states.

**Statement of significance:** Circadian rhythm, which is one of the most important biological rhythm, regulates and intervenes various cellular processes. Significant changes in the rhythmic dynamics may lead to pathological states which may trigger various diseases. In this work, the impact of CK2 via per gene mutants on rhythmic dynamics is investigated, and found three distinct states, namely, *active, weak activity* and *rhythmic death* driven by CK2 which may correspond to various cellular states. Noise due to intrinsic random molecular events and cellular size variability is found to have the capability of regulating and controlling rhythmic properties, and can trigger to the three rhythmic states. We then listed various possible pathways which are regulated by per gene mutants and corresponding various possible pathological states.

---

Life is a manifestation of rhythms. Circadian rhythm is one of such important rhythms, and is reported to be as an endogenous biological process which is entrainable oscillation of roughly 24 hours time period [1, 2]. This rhythm has been reported to be associated with wide range of living systems such as plants, animals, fungi, and cyanobacteria [4–6, 6]. Even though these rhythms are endogenous (i.e. self-sustained), they can be adjusted (entrained) to the local environment by external factors known as zeitgebers, for example, daylight, temperature, and many other molecular fluctuations [8, 9].

The oscillation in circadian rhythm is reported to be due to negative feedback loop in the genetic regulation which involves various clock proteins and genes in the biochemical reaction model [10, 11]. Further, it is reported that this rhythm is affected by various environmental factors, such as changes in sleeping conditions, various changes in the regulations of hormones such as testosterone, progesterone etc, variations in sunlight and various other factors. It is also reported that changes in the rhythmic process may lead to various disease conditions, such as, jet lag, cancer etc in the organism [8, 12].

The protein kinase CK2 (casein kinase II) is an important enzyme known as the heart of self-sustaining circadian clocks in animals, plants, fungi etc[13, 14]. CK2 is ubiquitous eukaryotic protein kinase present in both the nucleus and cytoplasm [15, 16]. In Drosophila, this CK2 phosphorylates and destabilizes the PERIOD (PER) and TIMELESS (TIM) proteins, which then inhibits CLOCK (CLK) transcriptional activity [17–20]. CK2 also targets the CLK activator directly. It is traditionally classified as a messenger-independent protein serine/threonine kinase and is typically found in tetrameric complexes consisting of two catalytic (alpha and/or alpha’) subunits and two regulatory beta subunits [21–23]. Further, CK2 has a vast array of candidate physiological targets and participates in a complex series of cellular functions, including the maintenance of cell viability [24–27]. Moreover, it is also reported that CK2 participates in cell cycle control, DNA repair, regulation of the circadian rhythm and other cellular processes. However, the regulating mechanism of CK2 to circadian rhythm pathway and its dynamics is still not fully studied.

Further, it is also reported that CK2 phosphorylates Per and Tim in vitro which indicates effect of CK2 upon Per and Tim protein [28–31]. Moreover, it is reported that Tim (amino acids 11159) protein is phosphorylated by CK2 to a lesser extent than Per. These data are consistent with the direct regulation of Per and Tim by CK2. Several reports have been published regarding role of Ck2 on drosophila circadian rhythm via Per protein direct interaction. However, It is still not clear that what is the exact mechanism by which it affects circadian rhythms at molecular level. However, the dynamics of clock molecular variables modulated by CK2 is still a debatable area of research, and need to be studied systematically. In the present study, we focussed on the stochastic approach to study the dynamical behavior of circadian rhythm driven by CK2 to understand the role of CK2 in regulating circadian rhythm at molecular level.

In this work, we extend the biochemical pathway model of drosophila by incorporating various possibilities of interaction of CK2 with clock proteins. The constructed model is presented in the materials and methods including method to simulate the biochemical network model in section 2. In section 3, we describ the numerical simulation results and interpretation. Next, we conclude our work based on the results obtained.

## Materials and methods

We propose an extended model of drosophila circadian model pathway which incorporates the impact of CK2 and describe the stochastic simulation algorithm used for the simulation of the biochemical reaction network as in the following.

### Description of circadian-CK2 model

The proposed *circadian-CK2* integrative model is an extension of the drosophila circadian rhythm model by Goldbeter [6] and Gonze et.al. [7] by incorporating the CK2 molecular pathway in the model based on some experimental reports which indicated interaction of CK2 protein with clock proteins in circadian rhythm [17, 18]. The model is shown in Fig. 1, and brief description of it is given as follows. This model is based on the repression exerted by the nuclear form of a clock protein (*P*_*N*_) on the transcription of its gene into mRNA (*M*_*P*_) [25]. mRNA is synthesized in the nucleus and transferred to the cytosol, where it accumulates at a maximum rate *k*_9_; there it is ubiquitinated by enzyme with a rate *k*_10_. The rate of synthesis of the protein *P*_*O*_ is proportional to the creation of *M*_*P*_ and is characterized by an apparent first order rate constant *k*_12_. Parameters *k*_14_ and *k*_19_ denote the maximum rate(s) and Michaelis constant(s) of the kinase and phosphatase involved in the reversible phosphorylation of *P*_*O*_ into *P*_1_ and *P*_1_ into *P*_*O*_, respectively. Parameters *k*_20_ and *k*_21_ denote the maximum rate(s) and Michaelis constant(s) of the kinase and phosphatase involved in the reversible phosphorylation of *P*_1_ into *P*_2_ and *P*_2_ into *P*_1_, respectively. The fully phosphorylated form *P*_2_ is degraded by an enzyme and is transported into the nucleus at a rate characterized by the apparent first-order rate constant *k*_29_. Transport of the nuclear form of the clock protein (*P*_*N*_) into the cytosol is characterized by the apparent first-order rate constant *k*_30_. It is reported that the negative feedback is exerted by the nuclear clock protein on gene transcription. CK2 phosphorylate the clock protein which may form three complex (*CK*2 − *P*_*O*_, *CK*2 − *P*_1_, and *CK*2 − *P*_2_) due to three different form(s) of the available proteins. Synthesis of *CK*2 − *P*_*O*_ is assumed to occur with a rate constant *k*_7_ and subsequently the dissociation of this complex is assumed to occur with rate constants *k*_8_. Similarly, the synthesis of CK2-*P*_1_ is considered to occur with a rate constant *k*_4_ and subsequently the dissociation of this complex is assumed to occur with rate constants *k*_5_. Further, synthesis of *CK*2 − *P*_2_ is taken to occur with a rate constant *k*_9_ and subsequently the dissociation of this complex is assumed to occur with rate constants *k*_10_. CK2 is ubiquitinated at rate constant *k*_6_. The synthesis of the CK2 in the network is assumed to occur at the rate constant *k*_3_. The CK2 kinase has a key function in the clockwork of various organisms. We describ list of all the molecular species related to the integrated model in Table 1. The chemical reactions, propensity function and their rate constants are listed in Table 2.

**Table 1.** List of molecular species

| S.No. | Molecular Species | Description |
| --- | --- | --- |
| 1. | $P_N$ | Nuclear Protein |
| 2. | $G$ | Promoter of the gene without ligand |
| 3. | $GP_N$ | G and $P_N$ Complex |
| 4. | $GP_{N2}$ | G and $P_N$ |
| 5. | $GP_{N3}$ | G and $P_N$ |
| 6. | $GP_{N4}$ | G and $P_N$ |
| 7. | $M_P$ | $mRNA$ |
| 8. | $E_m$ | Enzyme |
| 9. | $C_m$ | Complex |
| 10. | $P_0$ | Unphosphorylated Per protein |
| 11. | $E_1$ | Enzyme |
| 12. | $C_1$ | Complex |
| 13. | $P_1$ | De-phosphorylated Per protein |
| 14. | $E_2$ | Enzyme |
| 15. | $E_3$ | Enzyme |
| 16. | $C_2$ | Complex |
| 17. | $C_3$ | Complex |
| 18. | $P_2$ | Phosphorylated Per protein |
| 19. | $E_4$ | Enzyme |
| 20. | $C_4$ | Complex |
| 21. | $E_d$ | Enzyme |
| 22. | $E_d$ | Complex |
| 23. | $CK2$ | CK2 kinase enzyme |
| 24. | $CK2-P_0/CK2-P_1/CK2-P_2$ | Complex of CK2 and Per proteins<br>($P_0/P_1/P_2$ respectively) |

**Table 2.**
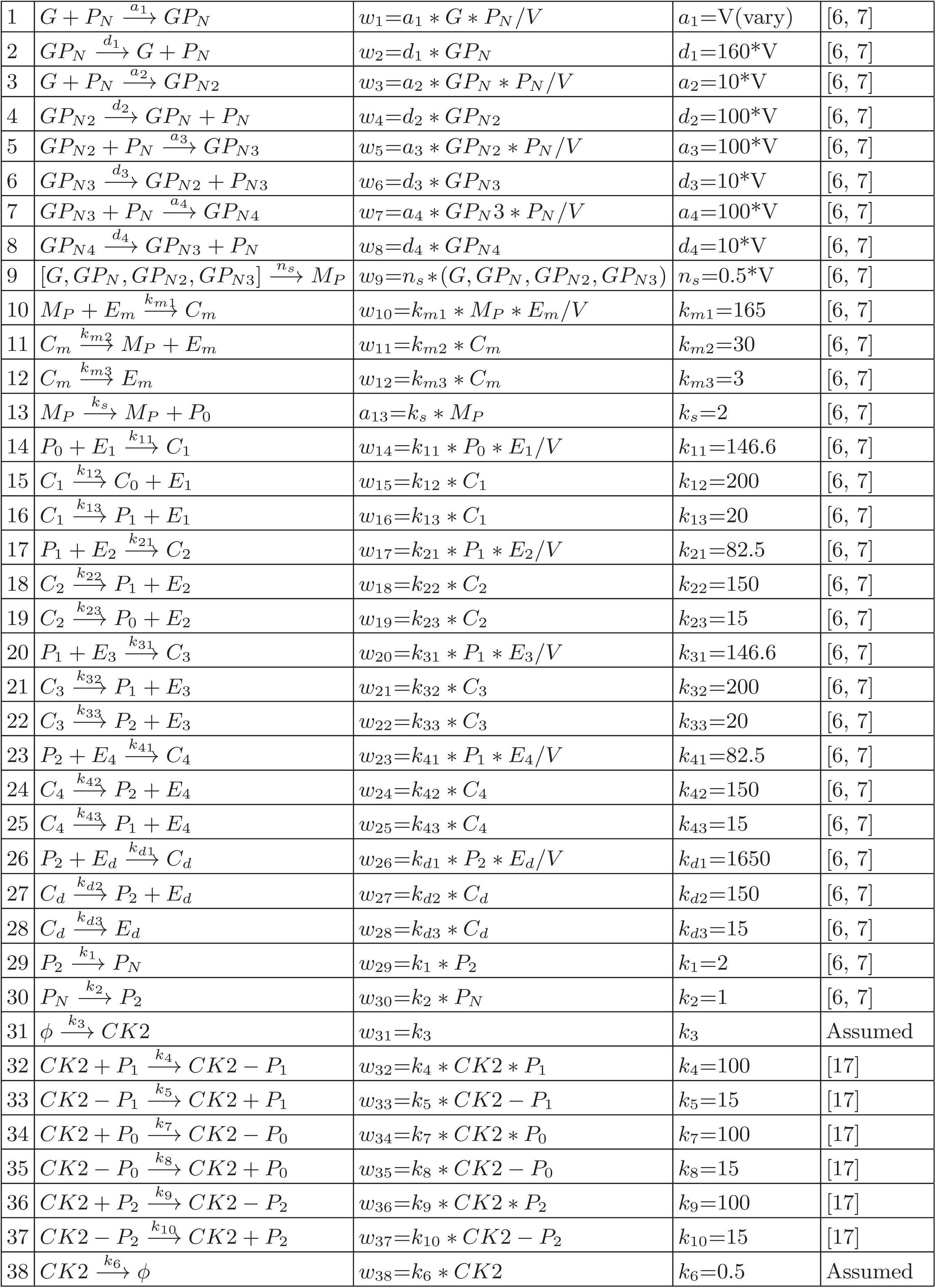
List of chemical reaction, propensity function and their rate constant

**FIG. 1:**
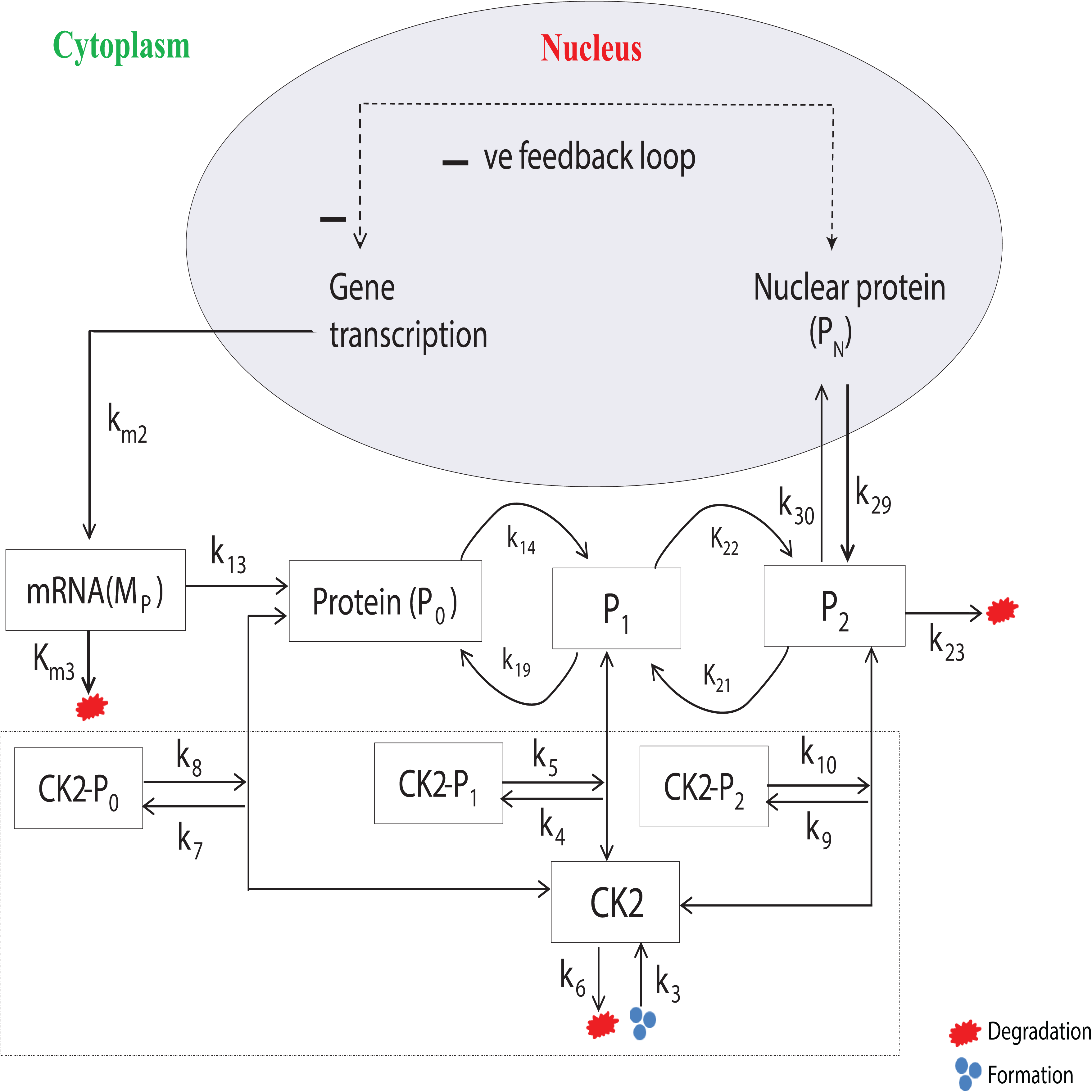
Schematic diagram of circadian rhythm driven by CK2.

### Simulation technique of biochemical reaction pathway

Complex dynamical processes governed by a set of well defined reaction channels are generally noise induced stochastic process due to random molecular interaction in the system (origin of *intrinsic noise*) and continuous interaction of the system with the random environmental fluctuations (origin of *extrinsic noise*) [32, 36, 37]. Consider the system has *N* molecular species variables, whose population vector is defined by, 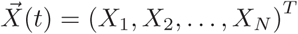, where *T* is the transpose of the vector, which undergo *M* reaction channels, *R*_*i*_, *i* = 1, 2, …, *M* given by,

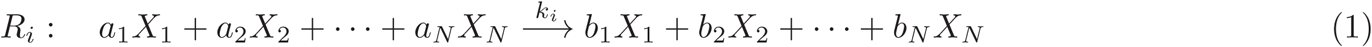

where, {*k*_*i*_}; *i* = 1, 2, …, *M* is the set of classical rate constants. The stochastic rate constant *c*_*i*_ of any its reaction can be expressed in terms of classical rate constant *k*_*i*_ by *c*_*i*_ = *k*_*i*_*V* ^1−*ν*^, where, *V* is the system size and *ν* is the state change parameter of the its reaction [32]. The trajectories of the variables provided by birth and death process due to molecular interaction given by equation (1) can be traced by solving Master equation, where the rate of change of configurational probability 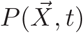 as a function of time can be described by,

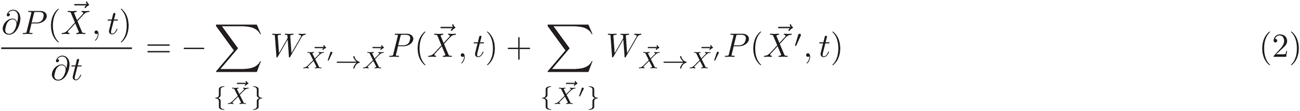

where, *W* and *W*′ are the transition probabilities of the two configurational states 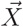 and 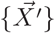. Except for simple system, solving master equation (2) analytically is very difficult. However, numerically one can solve the master equation (2) using stochastic simulation algorithm (SSA) or Doob-Gillespie algorithm due to Gillespie [32] based on the theoretical foundation developed by Doob JL [33, 34] and originally proposed by Kendall [35]. It is Monte carlo type algorithm which provides the exact numerical calculation by taking every possible interaction in the system [32]. This algorithm is in fact non spatial individual based analog of master equation (2) which is constructed on the physical basis of molecular collision in each reaction channel at certain constant temperature. This SSA is based on two important random processes, namely, reaction fire and reaction time which are two independent random processes. These two processes are served in this algorithm by generating two statistically independent random numbers *r*_1_ and *r*_2_ such that reaction time is computed using 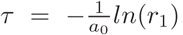, where, 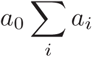, *a*_*i*_ is the *i*^*th*^ propensity function given by *a*_*i*_ = *h*_*i*_*c*_*i*_, where, *h*_*i*_ is the number of possible molecular combinations of ith reaction; and *k*^*th*^ reaction will fire when it satisfy, 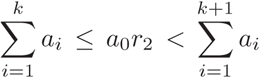. Intrinsic noise (*ξ*) associated with the species dynamics in the system is inversely proportional to the square root of the systems size *V* (i.e. 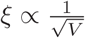) [38].

### Algorithm to calculate permutation entropy: Bandt and Pompe approach

Permutation entropy can be used to measure the complexity of a system associated with the dynamics of system’s variables [39, 40]. The basic algorithm of calculating permutation entropy *H* of a time series is as follows. Consider a dynamical variable *x*(*t*) of a system given by the discrete time series *x* = {*x*_*i*_} where *i* = 1, 2, …, *N*, and *N* is finite and total number of discrete time elements in the time series data. We define an embedding dimension *d*, preferably *d* = 3, …, 7 in order to represent the data in consecutive patterns of size of dimension *d*. For a particular value of *d*, there will be *M* possible permutated sequences of inequalities of sequence elements. If we take *d* = 3, we will have *M* = 6 arrangements of permutations given by, *u*_1_ = {*x*_1_, *x*_2_, *x*_3_}, *u*_2_ = {*x*_1_, *x*_3_, *x*_2_}, *u*_3_ = {*x*_2_, *x*_1_, *x*_3_}, *u*_4_ = {*x*_2_, *x*_3_, *x*_1_}, *u*_5_ = {*x*_3_, *x*_1_, *x*_2_}, *u*_6_ = {*x*_3_, *x*_2_, *x*_1_}, where, *x*_1_ ≠ *x*_2_ ≠ *x*_3_. Then we calculate the probability *p*_*u*_ for each single permutation sequence, which is the ratio of number of values for the particular sequence to the total number of all possible permutations for the embedding dimension *d* in the time series data. Now, we can calculate the simple Shannon entropy for a sequence by,

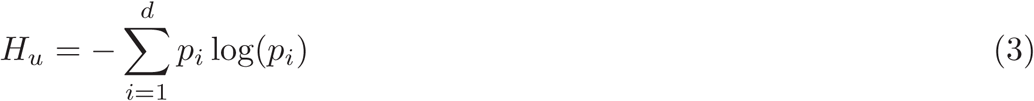

and the permutation entropy of embedded dimension *d* is given by the sum of these entropies as follows,

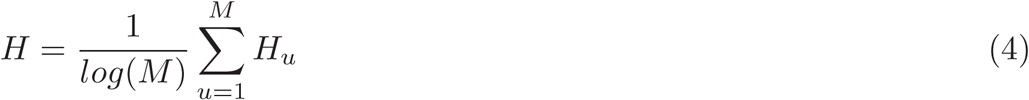

where, 0 ≤ *H* ≤ 1. The mapped permutation entropy spectrum of time series *x*(*t*) is indicated by *H* = {*H*_1_, *H*_2_, …, *H*_*N*_}, and is quite similar to the behavior of Lyapunov spectrum of the same time series [39].

## Results and discussion

We have done large scale numerical simulations of the proposed *circadian-CK2* integrated model were done using SSA [32] and results are being presented. Our main focus in this work would be to show the regulation of CK2 to the dynamics of clock proteins involved in circadian rhythm. We also present the impact of the noise upon the system variables is also be presented [41–43].

### Modulation of clock proteins (*P*_*N*_, *M*_*P*_) by CK2

We first present the results of how CK2 regulates clock protein (*P*_*N*_, *M*_*P*_) dynamics. Since population of CK2 protein in the system of size *V* (keeping *V* = 200 to be fixed) is proportional to the rate of creation of this protein in the system (*CK*2 ∝ *k*_3_), the variation in CK2 might cause change in the interaction rate of the other molecular species in the reaction network model. Hence, we look for the variation in the dynamics of the clock proteins driven by CK2 via change in the values of *k*_3_. First we allow all three phosphorylation of all cytosolic per proteins, *P*_0_, *P*_1_, *P*_2_ with CK2 to take place (Fig. 1), and found that for small *k*_3_ = 0.001 prominent oscillation (active state) in the in nuclear per protein *P*_*N*_ is exhibited with time period of oscillation, 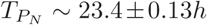 and amplitude of oscillation 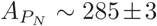 (Fig. 2 A upper panel). Further increase in *k*_3_ suppress the oscillation (*k*_3_ = 0.04, 0.08) allowing to increase *T*_*P*_*N* and decreasing *A*_*P*_*N* significantly which could correspond to weak circadian activity [44]. This increase in time period is due to increase in phosphorylation due to increased in interaction of CK2 with per protein (*P*_0_, *P*_1_, *P*_2_) as evident from the experimental reports of Albrecht et al. [45]. Further, the weak circadian activity may cause various diseases such as aging of brain, metabolic dysfunction, dementia, and many other cancer related diseases [46]. When the *k*_3_ is sufficiently large *k*_3_ ∼ 1.0 the *P*_*N*_ dynamics show both oscillation and amplitude death scenario [47]. This state may correspond to circadian rhythmic death or death of the living system [44, 46]. These three important states in the dynamics of *P*_*N*_ of the circadian rhythm induced by CK2 can be shown in two dimensional parameter spaces 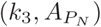 and 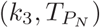 in Fig. 3. Here, the parameters 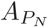 and 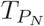 are mean values *P*_*N*_ amplitude and time period for a range of *t* = [10−500]*hrs*. In these phase diagram like plots all the three circadian states can be easily demarcated.

**FIG. 2:**
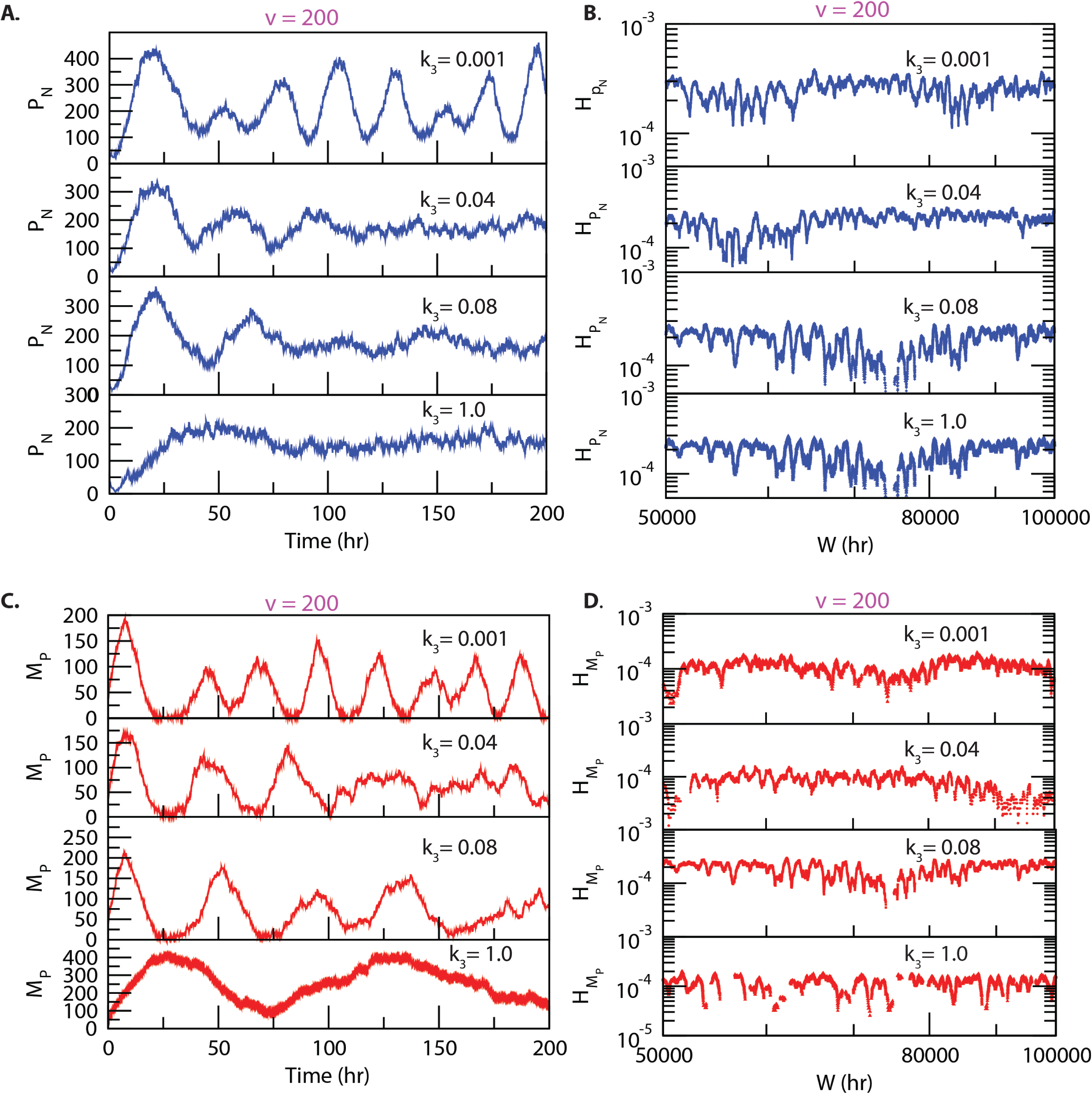
Dynamics of variables in circadian rhythm driven by CK2: We consider all possible interaction of CK2 with *P*_0_, *P*_1_ and *P*_2_ as shown in Fig. 1. A. Plots of dynamics of *P*_*N*_ for four various values of *k*3 for fixed value of system’s size *V* = 200. B. Permutation entropy spectrum, plots of 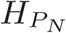 as a function of *Whr* for the corresponding time series on *P*_*N*_. C. Similar dynamics of *M*_*P*_for the same parameter values as in the case of *P*_*N*_, and D. Corresponding permutation entropy spectra corresponding to *M*_*P*_ dynamics.

**FIG. 3:**
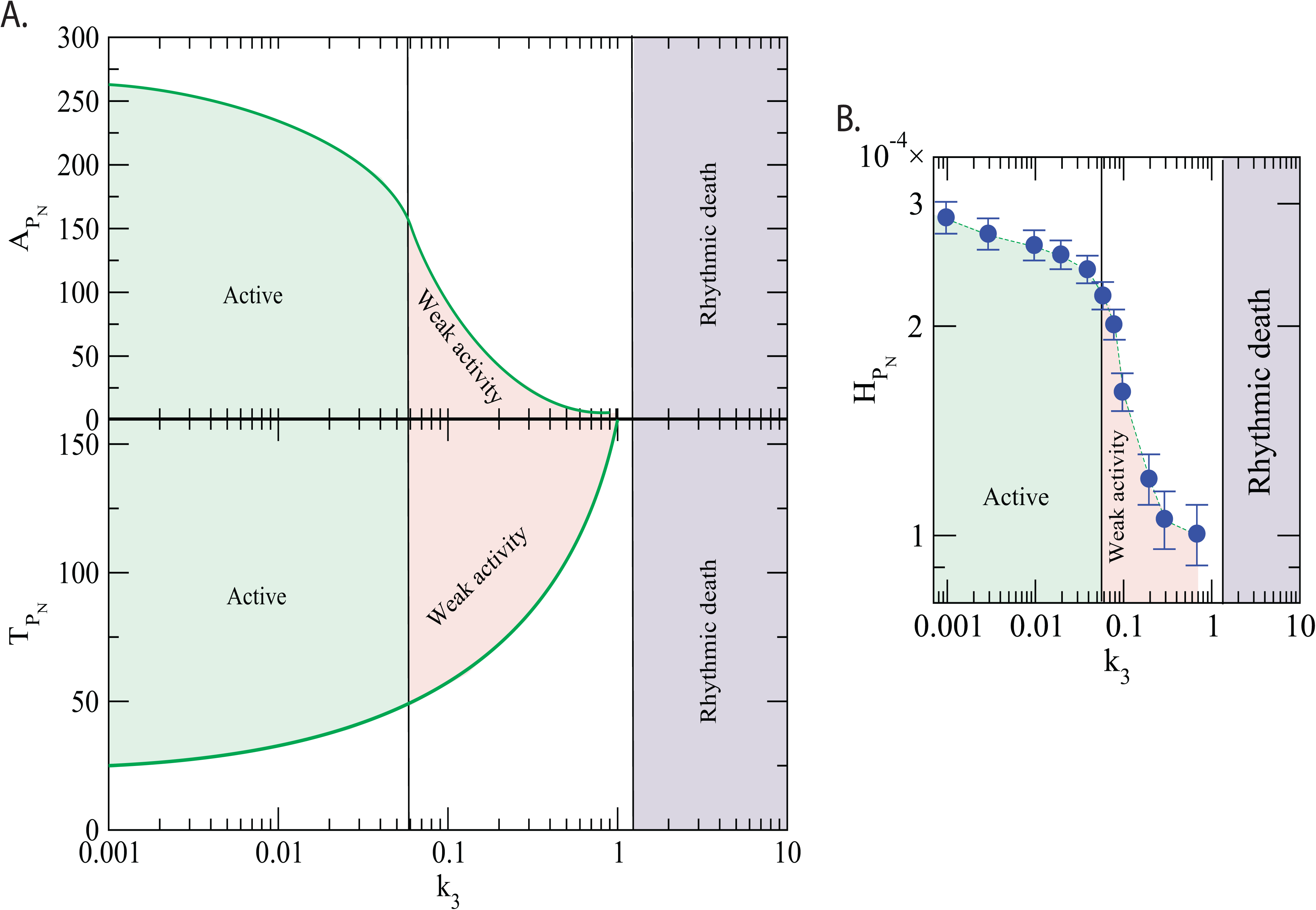
Circadian rhythm states driven by CK2: A. Plots of 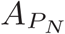 and 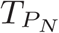 as a function of *k*_3_. Each point is in the curves are the average of the amplitude and time period in each time series between [10-200]hrs corresponding to single value of *k*_3_. Based on the behavior three different circadian states are identified, namely, *active, weak activity* and *rhythmic death*. B. Permutation entropy curves as a function of *k*3 with error bars and the three circadian states are demarcated.

We now study the measure of complexity in the three states obtained (active, weak activity and rhythmic death) driven by CK2 by calculating permutation entropy 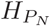 of the dynamics of *P*_*N*_ at the three respective states (Fig. 2 B). If we take average permutation entropies of active, weak activity and rhythmic death as 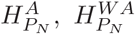 and 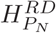, then our simulation results show that 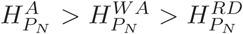. The three circadian rhythm states (active, weak activity, rhythmic death) can easily be classified using 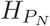(Fig. 3 right panel). In this phase like diagram, each point is the average of permutation spectrum for each value of *k*_3_ with error bars. Hence, the results indicate that 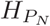 can be used as an order parameter to classify various states of the circadian rhythm. We propose that 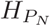 can be useful to clinicians and medical practitioners.

Similar nature is obtained in the dynamics of *M*_*P*_ as we have got in the case of *P*_*N*_ dynamics as shown in Fig. 2 C. In the same way, we also get similar behavior in permutation entropies at the three states (Fig. 2 D). These result suggests that CK2 plays a very dynamic role in the regulation of circadian rhythm in living systems. The change in rhythmic properties may cause various changes in the physiological processes of the organism that leads to various diseases to the organisms.

### Configurational impact of CK2 with per proteins on circadian rhythm

We now consider various possible configurational interaction of CK2 with any one of the per proteins *P*_0_, *P*_1_ and *P*_2_ to form complexes *CK*2 − *P*_0_, *CK*2 − *P*_1_ and *CK*2 − *P*_2_ respectively as shown in Fig. 4. All the simulations are done for the same values of *k*_3_ and *V* parameters as we did in the previous case where CK2 interacts with all *P*_0_, *P*_1_ and *P*_2_. The results show that if CK2 individually interact with any one of the per proteins via formation of their complex, *P*_*N*_ dynamics shows both active (for small values of *k*_3_) and weak activity (for larger values of *k*_3_) states (Fig.4). However, the system needs significantly large values of *k*_3_ to obtain rhythmic death state which is not shown in the figure. Further, configurational interaction of CK2 with *P*_2_ is much more sensitive in driving the circadian rhythm states as compared to the interaction of CK2 with *P*_1_ and *P*_0_ respectively. We then calculated 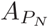 as well as 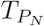 for each time series when CK2 is allowed to interaction with each one of *P*_0_, *P*_1_ and *P*_2_ in the *circadian-CK2* (Fig. 5 left panels). Each curve in 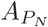 as well as in 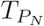, one can observe the three circadian rhythm states as shown in the Fig. 5. It is also found that interaction of CK2 with *P*_2_ is much sensitive than it’s interaction with either *P*_0_ or *P*_1_ because the three circadian states are found at smaller values of *k*_3_.

**FIG. 4:**
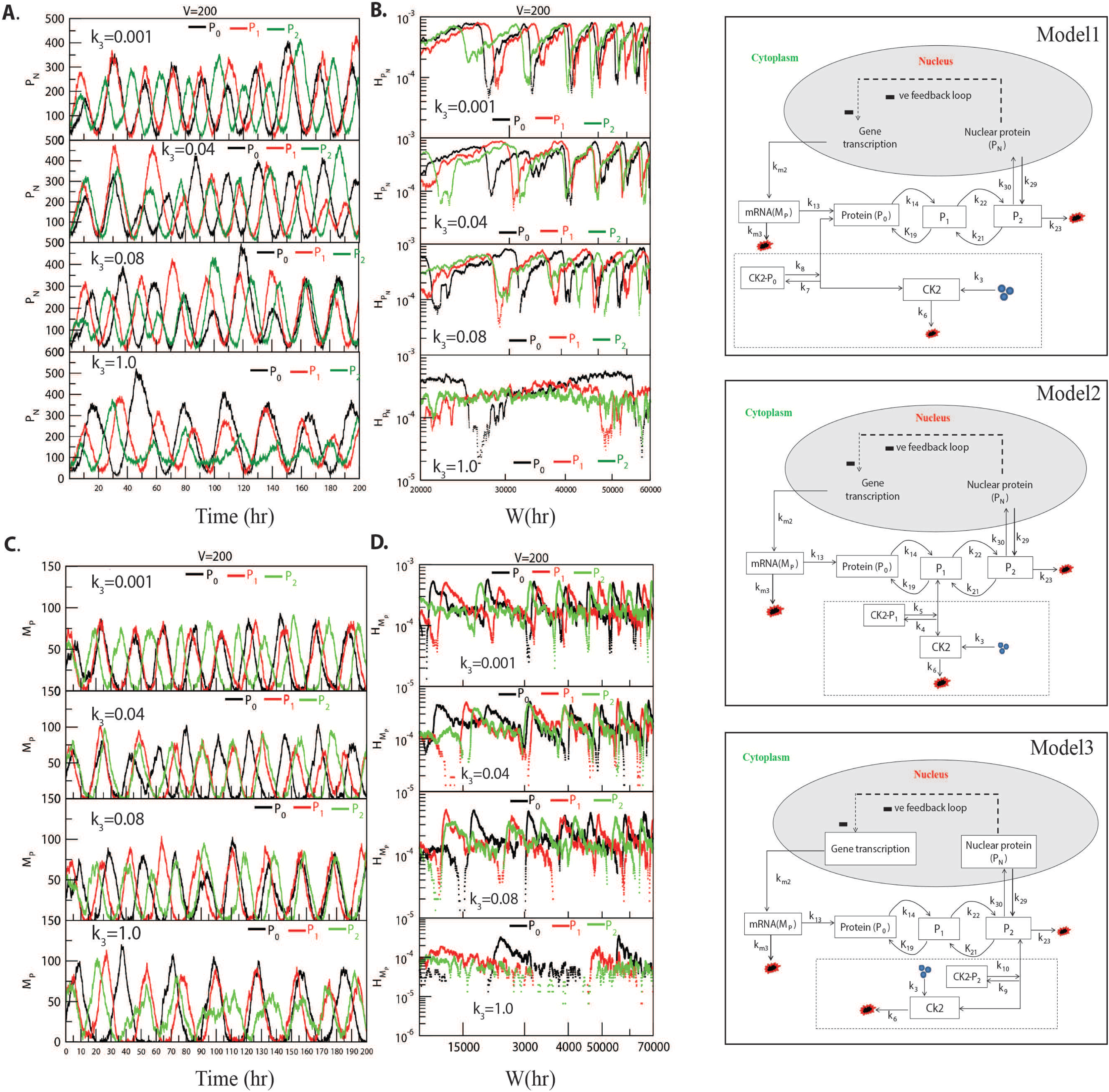
Different configurational interaction of CK2 with per gene mutants: A. Dynamics of *P*_*N*_ for different values of *k*3 when CK2 interacts with *P*_0_, *P*_1_ and *P*_2_; B. corresponding permutation entropy spectra (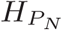 versus *W*). C. Dynamics of *MP* for the same parameter values as in *P*_*N*_; D. corresponding permutation entropy spectra for the same parameter values. Right hand column shows the models we have used for simulations.

**FIG. 5:**
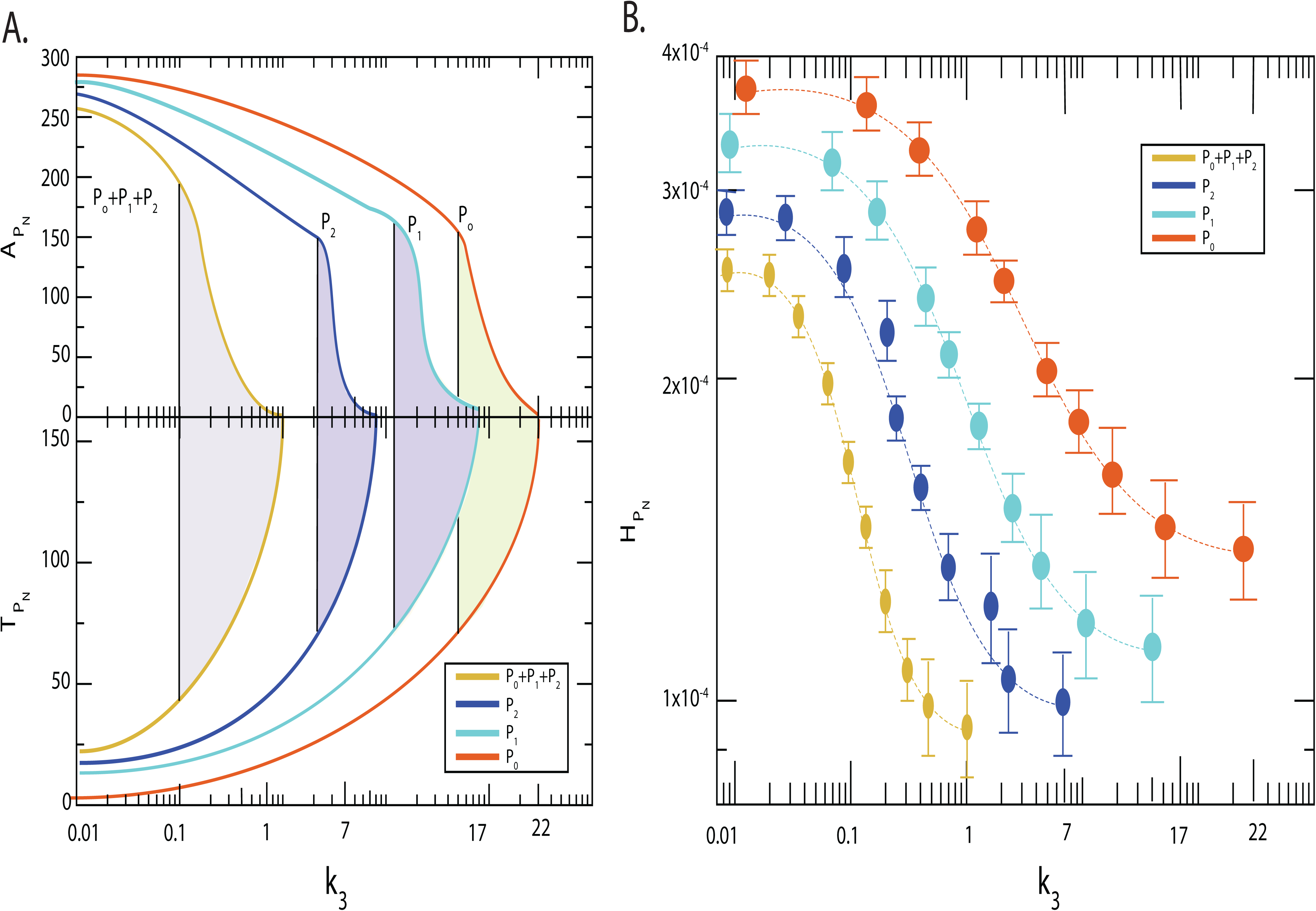
Phase diagram like behavior of circadian rhythm driven by CK2: A. Plots of 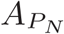 and 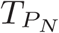 as a function of CK2 for all four configurational interaction of CK2 with *P*_0_, *P*_1_ and *P*_2_. B. Permutation entropy 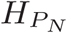 with respect to *k*_3_ for the four possible configurational interaction of CK2 with *P*_0_, *P*_1_ and *P*_2_.

The shaded colored areas in Fig. 5A are the regime of weak activity regimes when CK2 configurationally interact with any one of the per proteins (*P*_0_, *P*_1_, *P*_2_), which can also be known as *pathological states* of the corresponding configurations. The reason is that this is the regime where the properties of the rhythm, namely, amplitude and time period are significantly changed due to drastic increase in phosphorylation per protein/s with CK2 [45]. This drastic changed in circadian rhythm may cause various diseases starting from socio-psychological diseases [48], metabolic syndrome [49] to various types of cancer [50]. Excess phosphorylation of CK2 with per protein switch the pathological state to rhythmic death regime which could be the signature of apoptosis.

We then calculated permutation entropy 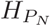 of the time series of *P*_*N*_ as a function of *k*_3_ for individual interaction of CK2 with *P*_0_, *P*_1_, and *P*_2_ respectively (Fig. 5 right panel). It shows that the three circadian states can be detected distinctly for each time series as in the previous section, and 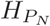 can be used as an order parameter to detect these states which can be useful to clinicians and medical practitioners.

### Noise can regulate the circadian states

Noise is everywhere, and is an inbuilt inherent property associated with the dynamics of any natural system [51]. We studied the dynamics of *P*_*N*_ for four possible configurational interaction of CK2 with the cytosolic per proteins (*P*_0_ + *P*_1_ + *P*_2_, *P*_0_, *P*_1_, and *P*_2_) as we did in the previous section by keeping *k*_3_ fixed (*k*_3_ = 0.01 which is in the regime of active circadian rhythm), and changing the strength of the noice 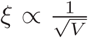 with four *V* values 80, 100, 200 and 500 respectively (Fig. 6 A). In all configurational interaction of CK2 with per protein, we observe that increase in noise (decrease in the value of *V*) can drive the *P*_*N*_ dynamics in the three circadian states, namely, active, weak activity and rhythmic death which we obtained in previous section. This indicates that noise can able to regulate the dynamics of circadian rhythm.

**FIG. 6:**
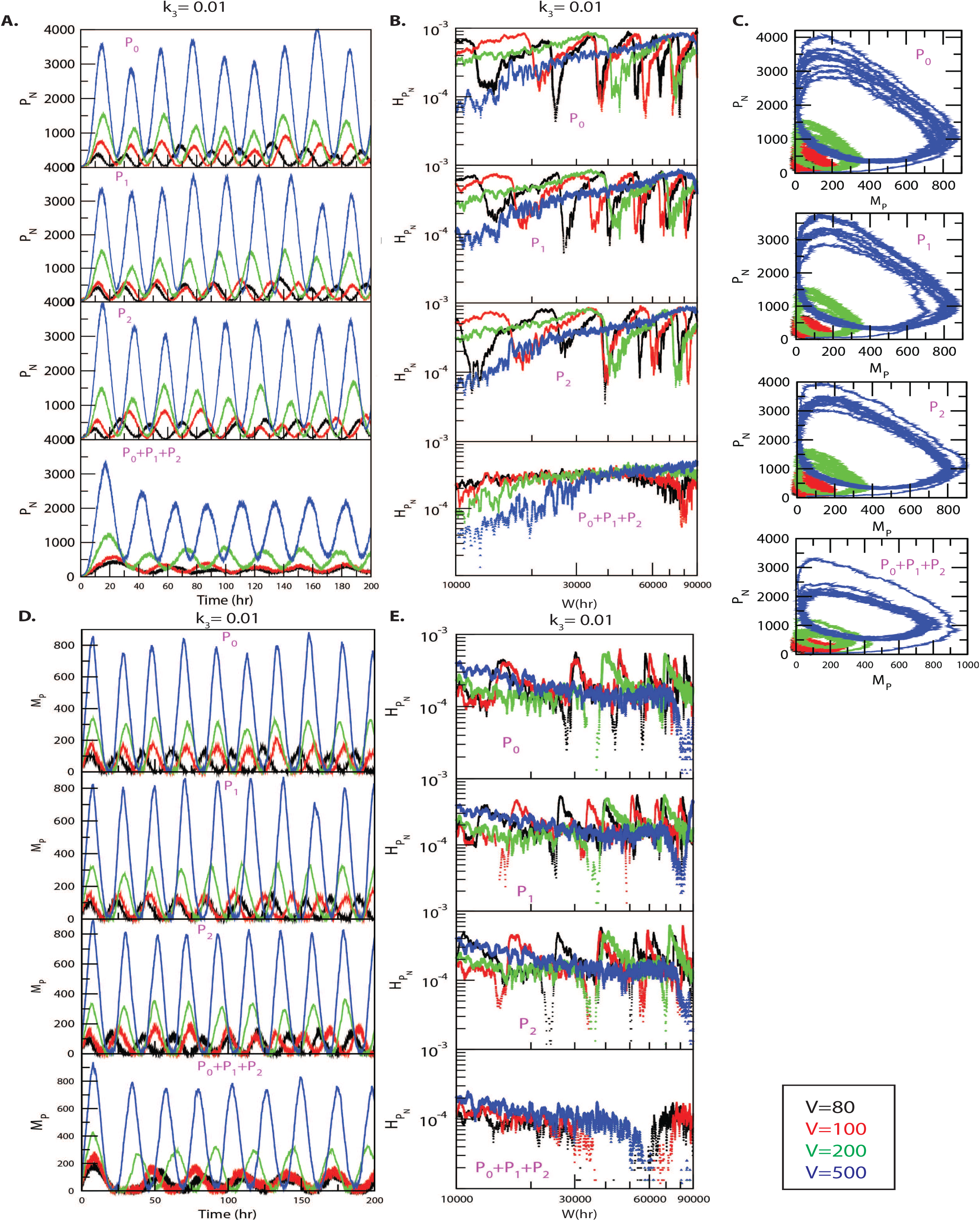
Noise induced *P*_*N*_ dynamics for all possible configurational interaction of CK2 with *P*_0_, *P*_1_ and *P*_2_: A. Dynamics of *P*_*N*_ for four different system sizes *V* = 80, 100, 200, 500 for fixed value of *k*_3_ = 0.01. B. Plots of 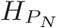 with respect to *W* for the corresponding system size values. C. The two dimensional plots of *P*_*N*_ and *M*_*P*_ for the four corresponding *V* values. D. Dynamics of *MP* for four values of *V*. E. Plots of 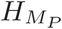 as a function of *W* for four *V* values.

Generally, the size of animal cell (both for proliferating and non-proliferating cells) can be changed due to various reasons [52], namely, cell growth and cell cycle progression [53, 54], starvation and pathological states in muscle cells [55, 56], dynamical cancer or tumor states [57, 58], defects in synaptic wiring/rewiring in neurons [59], manipulating extracellular signals to prevent apoptosis [60] etc. Animal cells can have cell size variability upto ∼ 59% of the normal cell size [60], and this change in the cell size can drastically affect the molecular crowding in the cell [61]. This variation in the system’s size is reflected as internal noise fluctuation (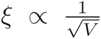 [38]) in the dynamics of the system’s variables. Since the circadian rhythm system is governed by well defines reaction channels (Table 2), this change in the molecular crowding could cause two main significant impacts in the system, first it allows to change the rate of interaction of the molecular species in the system. This change in molecular interaction leads to the change in the internal noise associated with the system dynamics reflected in the dynamics of the constituting variables *P*_*N*_, *M*_*P*_ and other species variables in the system. Hence, excess noise may destroy the signal associated with the system variables, and system may collapse. Second, this molecular crowing may trigger change in the molecular traffic in the system, where excess molecular crowding may affect traffic jam at which the system could not able to work in normal. This scenario could be the situation where the amplitude is minimized or death with infinitely large time period (Fig. 6 A). The similar behavior can be seed in the case of *M*_*P*_ dynamics (Fig. 6 D). The transition from fluctuated sustain oscillation (active state) to amplitude death (rhythmic death) via weak activity state can be seen in the two dimensional plot (*M*_*P*_, *P*_*N*_) as shown in Fig. 6 C.

Similar behavior can be observed in other three configurational interaction models of CK2 with *P*_0_, *P*_1_ and *P*_2_ respectively, where, noise is slightly more sensitive to the case where CK2 interacts with *P*_2_ than the other configurations (Fig. 6 A, C, D).

Now the amplitudes 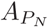 and time periods 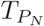 of *P*_*N*_ dynamics as a function of *V* are calculated for four different configurational interaction of CK2 with *P*_0_, *P*_1_ and *P*_2_ as shown in Fig. 7. From the results we can observe three distinct behaviors, namely, active, weak activity and rhythmic death driven by noise measured by *V* as marked in the figure. Hence, one can claim that noise in an important parameter which can trigger the system at various rhythmic states, and can able to regulate the system dynamics.

**FIG. 7:**
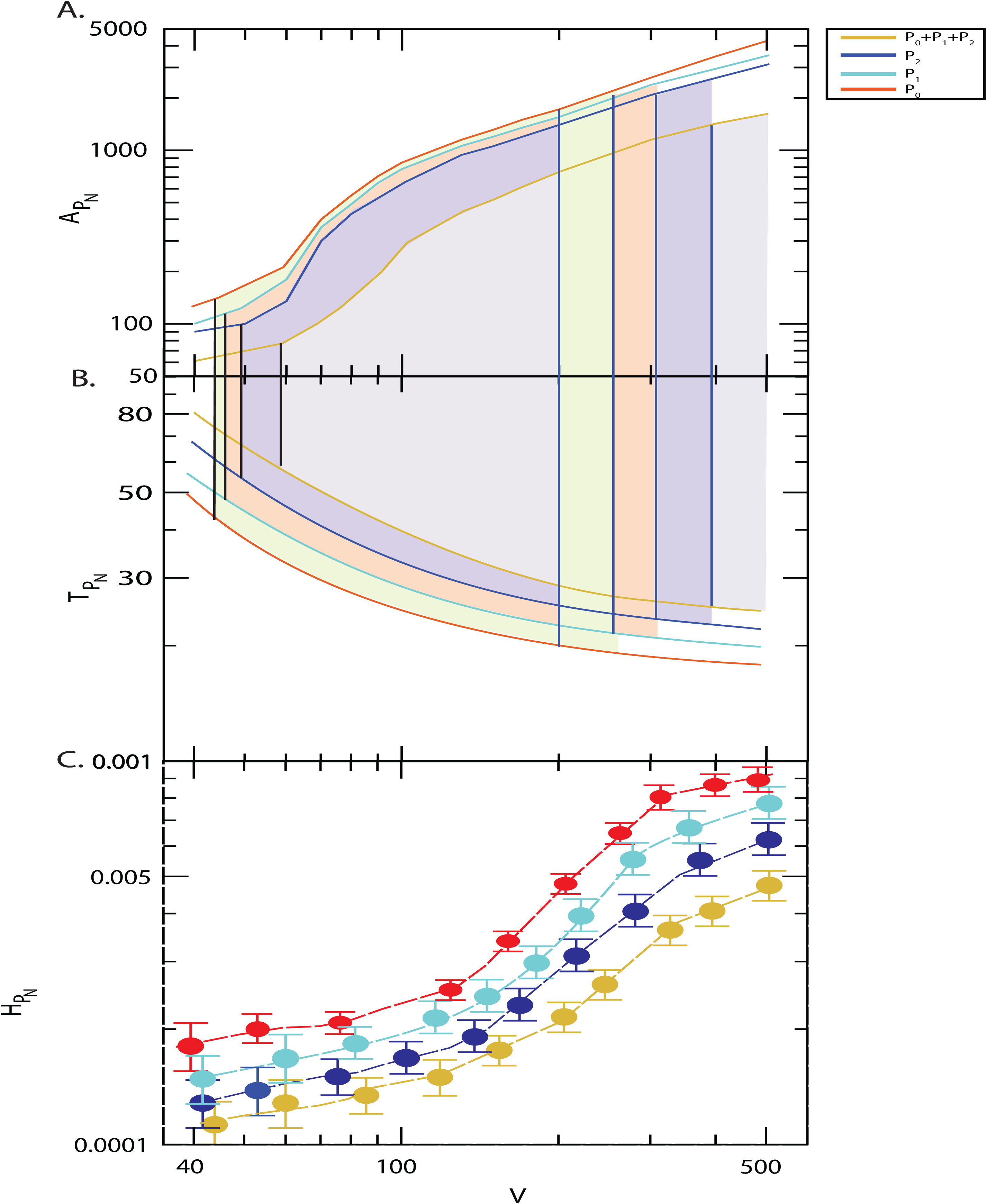
Noise induced circadian states: A. Plots of 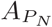 and 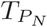 as a function of *V* for all four configurational interaction of CK2 with *P*_0_, *P*_1_ and *P*_2_ keeping *k*3 = 0.01. B. Permutation entropy 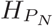 with respect to *V* for all four possible configurational interaction of CK2 with *P*_0_, *P*_1_ and *P*_2_.

Now we calculate the permutation entropy spectra of *P*_*N*_ and *M*_*P*_ corresponding to all the four models as a function of *V* values (Fig. 6 B and E). We also observe that the measure of 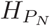 as a function of *V* can distinctly classify the three different circadian rhythm states (Fig. 7). If we denote 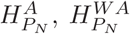, and 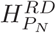 as permutation entropies corresponding to the circadian states active, weak activity and rhythmic death, then from the results we get that 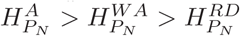 (Fig. 7).

### Circadian *per* induced cellular pathways which can trigger pathological states

We now search for various cellular pathways which can be triggered by per gene mutants (*P*_0_, *P*_1_, *P*_2_) in the circadian rhythm model as shown in Fig. 8. These cellular pathways can be grouped into three categories, first *cancer pathways* [50], *second socio-psychological pathways* [48, 62, 63], and *metabolic pathways* [49]. per genes with their own rhythms generally regulate other gene expressions participating various important cellular pathways whose significant changes may lead to various diseases. Because the disruption in the *P*_*N*_, *P*_0_, *P*_1_, *P*_2_ dynamics in the circadian rhythm may lead to drastic changes in the dynamics and mechanisms of these closely interacting pathways to various pathological states to cause various diseases corresponding to these pathways. Dysfunctional PER proteins disturb biological function like cell cycle, DNA damage, apoptosis, proliferation cause increase breast cancer, Breast cancer, Colorectal cancer, Prostate cancer, Lymphoma cancer, Lung Cancer, Liver cancer, Skin Cancer, Ovarian cancer, Brain cancer etc [64–68]. Metabolic functions are altered in PER cause increase Diabetes Obesity [71, 72]. PER protein aberration disrupts multiple biological functions such as aging Hydroxy methylation which cause increase Social physiological disease like Learning and memory impairment, Alzheimer’s disease [73]. The significant changes in the *P*_*N*_, *P*_0_, *P*_1_, *P*_2_ dynamics in the circadian rhythm may lead to drastic changes in the dynamics and mechanisms of these pathways to various pathological states to cause various diseases corresponding to these pathways. Hence, disruption in the *P*_*N*_, *P*_0_, *P*_1_, *P*_2_ dynamics by CK2 and noise in our study can trigger one or some of these pathways to pathological states leading to possible diseases corresponding to the pathways affected significantly. However, which pathway/pathways will be affected significantly by change in *P*_*N*_, *P*_0_, *P*_1_, *P*_2_ dynamics is still an open problem, and needs to investigate further.

**FIG. 8:**
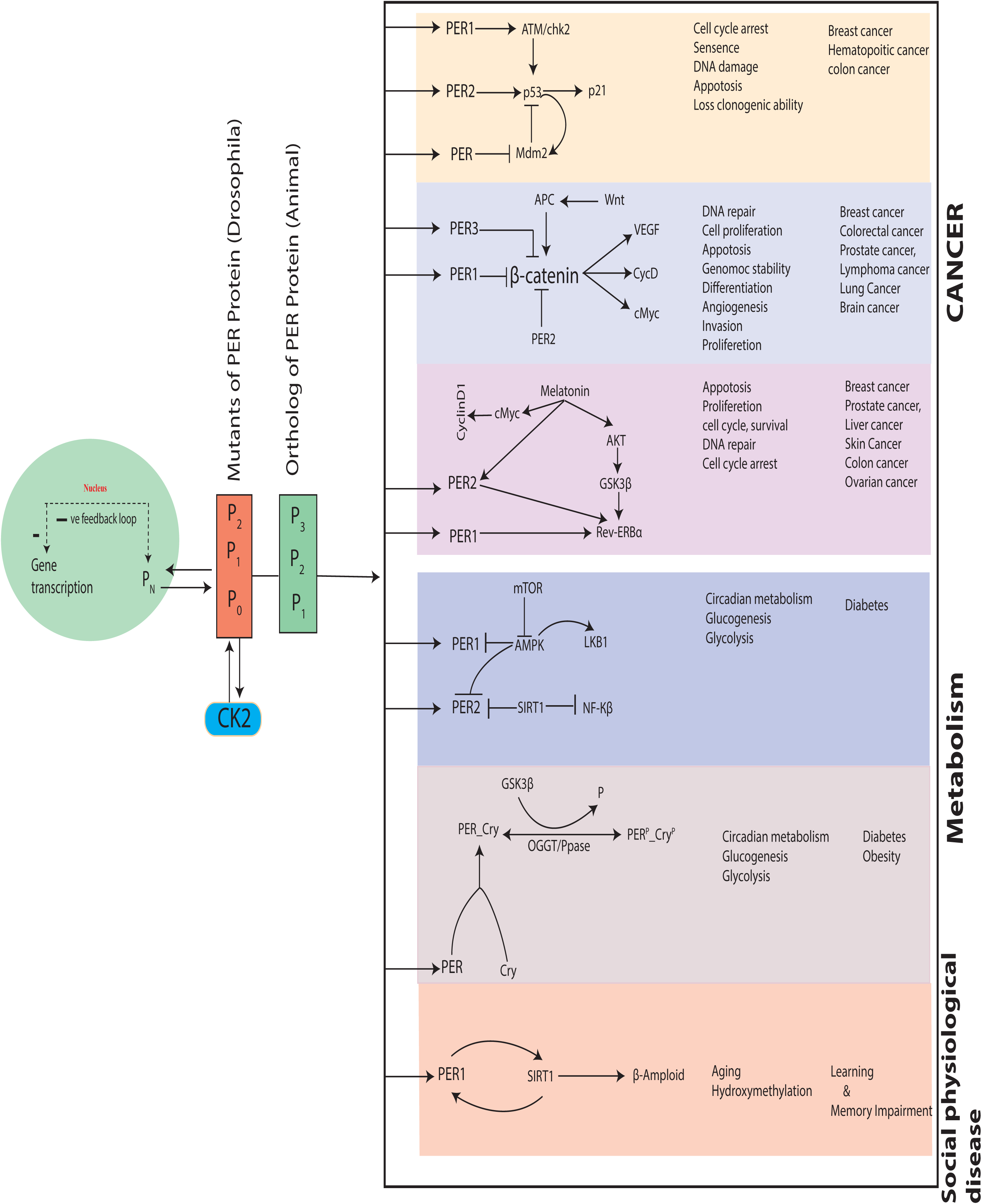
Possible pathological pathways driven by per gene variations: List of the pathological pathways which can be affected by variations in the rhythms of *per* gene mutants, and possible diseases.

On the other hand, patients having any one/some of the diseases mentioned in the Fig. 8 may become normal probably by monitoring proper circadian rhythm dynamics [65, 66]. The process can be termed as *reversed pathological state to normal*, which could be possible. However, even though it may be possible for some diseases, one needs to categorically study these possibilities from theoretical as well as experimental point of view. Hence, it has been proposed to study circadian rhythm genes/proteins as potential drug targets [69, 70].

## Conclusion

Circadian rhythm is one of the most important biological rhythms which can regulate and interfere various biological processes, such as, controlling brain activity, hormone production, cell regeneration and various other biological activities. In the present work we focussed on the impact of CK2 as well as stochastic noise on circadian rhythm model. It has been reported that CK2 promotes the progressive phosphorylation of clock protein which leads to the rapid degradation of hyperphosphorylated isoforms by the ubiquitin-proteasome pathway. Our results suggest CK2 availability in the systems has strong impact on the system’s dynamics reflected in the time evolution of *P*_*N*_ and *M*_*P*_. We found three distinct circadian rhythm states, namely, active, weak activity and rhythmic death driven by CK2. The active state corresponds to the time period of oscillation (*T*) in the rhythm variable (*P*_*N*_, *M*_*P*_, etc) is roughly of (22 − 24)hours with optimal amplitude *A*. If *T* is larger than that of at active state are termed as weak activity, whereas, the circadian state corresponding to *T* → ∞, where, *A* → 0 is known as rhythmic death.

Noise is an inherent property of any natural system associated with it. One interesting and important role of noise in a system is how it regulate the system and has the capability of controlling the system’s behavior. Since the size of animal cell can be changed due to various intracellular and extracellular functions [52], this variation in size is reflected as internal noise fluctuation in the system’s dynamics. In our study, we observe that noise can able to trigger the three circadian states (active, weak activity and rhythmic death), and can able to control system’s behavior. The modest nature of the biochemical defect supports the hypothesis that the circadian clock is highly sensitive to CK2 activity.

The radical change in the circadian rhythm could be due to various reasons, namely, genetic defects, shifting of works, aging etc [76]. This change in circadian state (i.e. at weak activity regime) may lead to various diseases specially cancer. The reason could be the variation in per gene mutants (*P*_0_, *P*_1_, *P*_2_(*drosophila*) → *P*_1_, *P*_2_, *P*_3_(*human*) [74]) may affect disruption in cell cycle through cMyc pathway with Per2, and ATK pathway with Per1 [76], and estrogen signal and HER with Per mutants in positive way which could be the signature of progression of breast cancer [75]. Further, since binding of Per to androgen receptor (AR) causes inhibition of AR transcriptional activity, the disruption of circadian rhythm may cause prostate cancer [77]. Not only these cancer types, this defects in circadian rhythm may cause various other types of cancer, namely, colorectal (due to Per2-ATM-Chk1/Chk2 pathway) [78], and many other types of cancer and tumorigenesis [50].

The disruption in circadian rhythm may also cause various other psychiatric and neurodegenerative diseases [48], jet lag and mental illness [62], sickness problems due to aging disorder [63], and many other diseases. This could be due to the fact that circadian rhythm pathway is associated with various important cellular pathways. Even some of the circadian genes/proteins are found to preserve cellular stability, for example, Per1 is experimentally found to be anti-apoptotic in nature [79]. Hence, one has to keep up with proper circadian rhythm in our day today life.

## Acknowledgments

M.Z.M. was financially supported by the Department of Health and Research, Ministry of Health and Family Welfare, Government of India under young scientist File No.R.12014/01/2018-HR, FTS No. 3146887. R.K.B.S. is financially supported by UPE-II, New Delhi, India, under sanction no. 101. This work is financially supported by University Grants Commission, New Delhi, India.

